# *lickcalc:* Easy analysis of lick microstructure in experiments of rodent ingestive behaviour

**DOI:** 10.64898/2026.03.09.710511

**Authors:** K. Linnea Volcko, James E. McCutcheon

## Abstract

Lick microstructure is a term used in behavioural neuroscience to describe the information that can be obtained from a detailed examination of rodent drinking behaviour. Rather than simply recording total intake (volume consumed), lick microstructure examines how licks are grouped, and the spacing of these groups of licks. This type of analysis can provide important insights into why an animal is drinking, for example, whether it is influenced by taste or affected by consequences of consumption (e.g., feeling “full”). Here we present a software package, *lickcalc*, that allows detailed microstructural analysis of licking patterns. The software is browser-based and is hosted at https://lickcalc.uit.no or the repository can be downloaded and installed locally. Lick timestamps can be loaded from a variety of formats and different analysis and plotting options allow quality control of data and determining critical parameters for microstructural analysis number and size of lick bursts. Data can be divided into epochs for detailed examination of changes across session. Batch processing and custom configurations are supported. In this manuscript, we demonstrate use of the functions exposed by *lickcalc* by analysing data comparing lick patterns between mice on a protein-restricted and control (non-restricted) diet. We show that lickcalc allows quality control of the data and uncovering of subtle differences in lick behaviour that are not apparent when just considering the total number of licks. This software makes microstructural analysis accessible to any researchers who wish to employ it while providing sophisticated analyses with high scientific value.

## Introduction

*lickcalc* is an open source software suite that performs microstructural lick analysis on timestamps of lick onsets and/or offsets. Microstructural analysis was first described in (Davis & Smith, 1992) and has since then been used to understand diverse phenomena. In-depth reviews on many of these, and microstructural parameters used to study them, are available (Johnson, 2018; Naneix et al., 2020; Smith, 2001). Briefly, lick microstructure can provide nuanced information about why an animal is drinking. By considering the distribution of interlick intervals (ILIs), individual licks can be grouped into bursts (also sometimes referred to as clusters or bouts) based on a minimum ILI that separates each burst. Outputs such as the average size of each burst and the total number of bursts can then be derived. The former output, burst size, is thought to vary with palatability of a solution and is suggested to reflect the affective state or hedonic experience associated with intake. As such, burst size increases with increasing concentrations of sucrose solution (Davis & Smith, 1992). The latter output, burst number, is proposed to represent either post-ingestive feedback (signals from the periphery are important for terminating meals and thus should reduce burst number) and/or motivation (when animals are more motivated to consume, burst number should increase). Concordant with this framework, burst number decreases with manipulations that mimic post-ingestive satiating feedback (e.g., CCK agonists; Overduin et al., 2014) and increases with manipulations that increase motivational drive (e.g., food restriction; Spector et al., 1998). Importantly, many manipulations will not affect both outputs equally so effects on palatability can be disentangled from effects on post-ingestive signalling.

Often, changes in microstructure are accompanied by changes in total intake, but this is not always the case: sometimes, equal intake will be achieved by quite different licking patterns that indicate changes in orosensory and post-ingestive feedback (Johnson et al., 2010; Volcko et al., 2020). Analysing lick microstructure is therefore highly valuable when trying to understand how a manipulation, *X*, affects appetite; if *X* causes an animal to feel more satiated after drinking, that may lead to a different interpretation than if *X* were to reduce the palatability of the solution.

Although much of the foundational work on drinking microstructure was on licking for nutritive solutions (e.g., sucrose solutions), microstructural analysis can also be used to study intake of water (McKay & Daniels, 2013; Santollo et al., 2021), ethanol (Patwell et al., 2021), and other tastants such as sodium, quinine, and non-caloric artificial sweeteners (J.-Y. Lin et al., 2012; Spector & St. John, 1998; Verharen et al., 2019). Lick microstructure has been used to shed light on, for example, how licking is affected by neuropeptides (McKay & Daniels, 2013), enzymes in the mouth (Chometton et al., 2022), ovarian hormones (Santollo et al., 2021), nutrient restriction (Naneix et al., 2020), responses to alcohol (Patwell et al., 2021), and diet manipulations (Johnson, 2012; Murphy et al., 2018; Volcko & McCutcheon, 2022).

Because of the value of microstructural data, many labs habitually record and analyse it. There are many others, however, that have not yet begun collecting and/or analysing these data. Investing in lickometers can be costly, but there are an increasing number of cost-effective alternatives to commercial products. As such, several open-source lickometer designs are now available (e.g. (Frie & Khokhar, 2024; Monfared et al., 2024; Petersen et al., 2024; Raymond et al., 2018; Silva et al., 2024).

Recording individual licks with high temporal resolution is necessary for microstructural analysis of drinking behaviour, but another barrier to reporting microstructure is its analysis. Here, we present an open source software solution, *lickcalc*, that allows lick microstructure to be analysed in an easy and intuitive manner. We include data from experiments in our lab illustrating benefits to using the software including quality control of data and gaining a deeper understanding of licking behaviour both with and without a difference in absolute intake.

## Materials and Methods

### Ethical statement

All animal care and experimentation complied with the EU directive 2010/63/EU for animal experiments. All experiments were approved by the Norwegian Food Safety Authority (FOTS protocol #22315).

### Getting started with *lickcalc*

*lickcalc* is hosted by UiT The Arctic University of Norway and can be accessed at https://lickcalc.uit.no. Alternatively, it can be installed locally following instructions in the repository (https://github.com/uit-no/lickcalc). To use *lickcalc*, the user drags a file into the application, changes file format if necessary, and indicates which column contains the lick onsets (and, if applicable, the lick offsets). Plots are automatically generated e.g., a histogram of licks across the session. Parameters such as session length can be manually changed in the interface or set in the optional configuration (config) file. For curious readers who do not have ready access to appropriate data, a selection of example files in different formats can be downloaded by clicking the “About” button in the top right corner of the app or at https://github.com/uit-no/lickcalc/tree/master/assets/examples. In addition, synthetic data files representing edge cases allow further validation for new users as they can observe the effect of changing parameters and producing expected effects and/or failures. When the university-hosted app is used, data will be transferred to the application instance on university servers for processing. Data are not explicitly stored but server infrastructure can still retain some data (e.g., web server logs, crash dumps etc). If users need strict local-only processing, then the repository can be cloned and the app installed locally.

### Microstructural analysis and quality control

Microstructural analysis is, in essence, a division of individual licks into groups of licks. To perform this grouping, the user must set several parameters. The first step is determining the threshold at which a burst is considered terminated. This is the **interlick interval (ILI) threshold**, which is the minimum amount of time licks must be separated by to be considered separate bursts. Early studies identified ILIs of 251-500 ms as separating “bursts” of licking, and pauses of >500 ms as separating “clusters” of licking (i.e., a cluster of licks is made up of several bursts of licking). Others have argued that an ILI threshold of 1 s provides better separation of lick bursts (Spector & St. John, 1998). In *lickcalc*, the user can set the ILI to any value (values between 250 ms and 3 s provided by default but can be adjusted using the config file). Another parameter that needs to be decided prior to the lick analysis is the **minimum licks per burst**. *lickcalc* allows between 1 and 5 licks by default. The appropriate number of for this parameter may vary depending on experimental set up, and the likelihood that a single lick represents a lick rather than, for example, a paw touching the spout. Finally, in *lickcalc*, the user can set a **long-lick threshold** between 0.1 and 1 s. This parameter is only available when lick offsets are included. Licks that are longer than the set threshold are counted as “long” and may indicate a problem (e.g., fluid bridges or an animal holding the spout with its paws) rather than a true lick. The user can decide whether to remove these long licks or not. Each of these parameters can be set manually or through the config file. Five plots are initially generated (see *Key Features* section below) and dropdown menus provide several additional plotting options. Tables are displayed showing values of several properties: total licks, intraburst frequency, lick length mode, intercontact mode, number of long licks, maximum long lick duration, number of bursts, mean licks per burst, mean interburst interval, Weibull alpha, Weibull beta, and Weibull r^2^.

### Saving and exporting data

To save these data, the user has two options. The first is to export a single Excel file in which the user sets the animal ID and chooses which data to export. These data allow the user to recreate the plots displayed in *lickcalc* or perform further analyses. The second option for saving the data is to add the loaded data to the *Results Summary* table. The results in this table remain even as new data files are loaded, so the data from many sessions (and/or individual animals) can be exported into a single Excel file. In addition to the data from the whole session, the user can choose to divide the session into epochs, or to examine only the first *n* bursts, or perform a trial-based analysis (e.g., for Davis rig experiments). Each of these analysis epochs can be added to the table. The table contains the data and the analysis parameters (e.g., minimum burst size) used to generate them. Finally, a batch process feature is available allowing multiple files to be analysed using the same parameters.

### Animals and housing

Male C57BL/6NRj mice (6-8 weeks old) were purchased from Janvier (France) and housed in a temperature- and humidity-controlled room. A 12:12 light cycle was used and all testing took place during the light phase. Mice from these experiments were run in three separate cohorts. Cohorts 1 (n=16) and 2 (n=16) were single-housed, due to prevalence of fighting in previous cohorts. Cohort 3 (n=16) was contact-housed, in which two mice shared a cage separated by a perforated divider that prevented physical contact while allowing visual, olfactory, and auditory contact between the cage-mates. Water and food were available ad libitum. In each cohort, half the mice were fed a control, non-restricted diet (NR; 20% casein; D11051801; Research Diets) and the other half an isocaloric protein-restricted diet (PR; 5% casein; D15100602; Research Diets). Mice had been on these diets for over two weeks before testing. Each cohort was used for a separate experiment before the experiment presented in this manuscript. Mice from cohort 1 were included in our previously published study (Volcko & McCutcheon, 2022) and as such had experience drinking 4% casein and 4% maltodextrin solutions in operant chambers and in their homecage. Mice from cohort 2 were used in an (unpublished) experiment in which they experienced drinking 4% maltodextrin and 4% collagen, both in operant chambers and their homecage. Mice in cohort 3 had experienced drinking 4% collagen, with or without additional tryptophan, in operant chambers as well as their homecage. Although the previous nutrient exposure differed between the mice in these cohorts, their experience with licking for nutritive solutions in the operant chambers was similar.

### Testing apparatus

Testing took place in operant chambers (24 cm x 20 cm; Med Associates, Fairfax, VT) equipped with a bottle connected to a contact lickometer. The sipper was accessible through a circular hole, 20 mm in diameter. The fan and house light were on continuously during testing. Hardware was controlled and the onset and offset of each lick were recorded with 1 ms precision using MEDPC-V software (Med Associates).

### Procedure

Mice in cohorts 1 and 2 were given two 1 h sessions in the operant chamber, with 1 kcal/ml of Resource Complete Neutral (9% kcal from fat, 23% kcal from protein, 68% kcal from carbohydrate; hereafter, “Ensure”) provided. The second session was used for the present analysis. Mice in cohort 3, on the other hand, received 4 days of homecage access to 1 kcal/ml Scandishake (43.7% kcal from fat, 3.8% kcal from protein, 52.7% kcal from carbohydrate). The solution was changed daily. A single 1 h session in the testing apparatus was then conducted. The primary purpose of these tests was to collect pilot data (e.g., to get a sense of how many licks mice in different diet conditions licked for these mixed-nutrient solutions).

### Data analysis and availability

The number and timing of licks was recorded. For microstructural analysis, we defined a burst as consisting of at least 3 licks with <1 s interlick intervals. Only mice that licked at least 100 times were included in the analysis. This meant that 2 NR mice from cohort 1 (with 2 and 9 licks), and 1 NR mouse from cohort 3 (52 licks), were excluded. For Weibull analysis, only mice with r^2^ > 0.8 were included. This resulted in 1 NR mouse being excluded. For statistical analysis between groups, two-tailed unpaired t-tests were used except for when the session was divided into epochs when a two-way repeated measures ANOVA was used (epoch as within-subjects and diet group as between-subjects factor. Statistical analysis was performed in Python using the packages *scipy* (Virtanen et al., 2020) and *pingouin* (Vallat, 2018) and code can be found in the repository below.

For users that want to install *lickcalc* locally, all necessary files are available at https://github.com/uit-no/lickcalc. The application makes use of the *lickcalc* function that is part of the *trompy* package and so using *lickcalc* without the graphical interface can be accomplished by just installing *trompy* (available via *pip* or at https://github.com/mccutcheonlab/trompy). All code and data used to produce this manuscript including Jupyter notebooks with figures are available at https://github.com/mccutcheonlab/lickcalc-paper.

This software – the *lickcalc* app and the underlying functions from *trompy* were created without the aid of AI coding tools, but for recent refinements (e.g., refactoring and organising the codebase and drafting the documentation), Github Copilot was used. No AI tools were used to prepare the manuscript. The authors take full responsibility for the contents.

## Results

*lickcalc* does not require any special software or coding knowledge: all the user has to do is load a file with timestamps of lick onsets (and, ideally, offsets) into the application and *lickcalc* will generate detailed microstructural analysis with a high degree of user control over key parameters. Resulting data provide values for number of licks, number of bursts, and burst size (among others) – the values that are often reported and used to draw inferences about post-ingestive and orosensory feedback of the solution. Importantly, several plots are produced that show information that helps with quality control and challenges the user to think critically about which parameters they have chosen.

### Key analysis outputs

Microstructural analysis with *lickcalc* produces several key outputs. We provide some information on each of these below for context, but the interested reader is referred to the cited literature for more in-depth discussion. For each of these outputs a mean is typically calculated but the full distribution is plotted and can also be exported.

**Intraburst lick frequency** is calculated from all ILIs within a burst of licking (i.e., those below the ILI threshold). While a rodent is licking, its tongue makes rhythmic protrusions that are under the control of a central pattern generator (Travers et al., 1997). Rats typically lick 6-7 times per second (Davis & Smith, 1992), while mice lick at a slightly higher rate (Johnson et al., 2010). In addition to these species differences, there are also strain differences (Johnson et al., 2010; St John et al., 2017). Because intraburst lick rate is under the control of the central pattern generator, it should remain relatively stable across mice and conditions (although recent studies have challenged this view and some manipulations might be expected to cause changes in this parameter) (X. B. Lin et al., 2013). A typical chart for a mouse might show a sharp peak around an intraburst ILI of ∼130 ms, which corresponds to a lick rate of 7.7 Hz. Much smaller peaks are often present at the harmonics of the intraburst ILI (e.g., a primary peak at 130 ms will have smaller peaks at 260 ms, 390 ms, and so on), often because of “missed licks” in which the animal attempts to lick but its tongue misses the spout. Many of these, or other differences from the expected pattern of results, may indicate problems with the experimental setup (e.g., if the animal fails to reach the spout frequently, then perhaps the spout is too far away) and so is a critical parameter for quality control. The intraburst ILI consists of time when the tongue is in contact with the spout and time when it is not. Thus, changes in intraburst frequency can be caused by alterations in lick length or the time when the tongue is not in contact with the spout (intercontact time; ILI - lick length). Examining each of these parameters will provide information about function and modulation of the central pattern generator driving licking behaviour. If lick offsets are available, then both these parameters are calculated (see below).

**Lick length** represents the time that the animal’s tongue is in contact with the spout and is only available when lick offsets are included in the input file. **Intercontact time** is the time in between licks when the tongue is not in contact with the spout. As with intraburst lick frequency, both these measurements should show little variability, and the graphs will have a sharp peak. Occasionally a lickometer will register longer licks than normal. A common cause of this is formation of a fluid bridge between the tongue and the spout during periods of high frequency licking. This can be prevented by moving the bottle further from the animal. In addition, other causes are if a rodent grabs the spout with its paws, or if a fluid droplet hangs between the spout and the cage and thus completes the electrical circuit. Concerns about data quality may be warranted with increasing number and duration of long licks. *lickcalc* displays both the number and maximum duration of licks above the threshold that the user has set. There is also an option to remove these problematic licks from the dataset.

**Burst frequency** describes how often certain burst sizes occurred. This is informative because burst size, by virtue of being a mean (mean licks per burst), does not consider potentially relevant information about the distribution of burst sizes. For example, a burst size of 80 could result from bursts all containing between 70 and 90 licks, or from many single licks and one or two bursts with hundreds of licks. The latter case might raise some questions about how reliable mean burst size is. Although single licks occur, they can also be caused by non-tongue contact with the lickometer. Changing the minimum licks per burst parameter can filter out some of these suspect “licks.”

#### Weibull probability

The Weibull analysis, as described in (Davis, 1996), uses a mathematical equation to fit burst size data to a survival function. Although used by some (Aja et al., 2001; Moran et al., 1998; Spector & St. John, 1998), it is still relatively rare to find Weibull probabilities in microstructural analyses. The Weibull function can be used on several aspects of data, such as lick rate across a session, but in *lickcalc* the Weibull probability is calculated for burst size. It plots the probability that, given *n* licks, the animal will continue to lick. This makes it sensitive to the licks per burst parameter that is set by the user. The Weibull α and β values reflect the slope (or decay) and shape parameters, respectively. Slope (α) has been shown to vary with palatability. One reason to use estimates from the Weibull distribution is that the mean licks per burst may not faithfully reflect the underlying burst distribution.

Next, we use data from our lab to demonstrate use of *lickcalc*, and the parameters arising from it, to perform quality control and uncover better understanding of the psychological and physiological drivers of ingestive behaviour.

### Using microstructural analysis to assess data quality and set burst parameters

The intraburst lick frequency can be examined to determine whether the majority of licks (which occur within bursts) conform to a tight distribution expected from a central pattern generator. In mice licking for “Ensure”, a sharp peak is seen at an ILI of 120-130 ms for both non-restricted (NR) and protein-restricted (PR) mice (Figs. 1A and 1B). This corresponds to a lick rate of approximately 7.7-8.3 Hz with no difference between groups (Fig. 1C; t = .217, p = .830). A smaller peak is apparent (especially in the restricted group) at double the peak ILI (the first harmonic), which represents licks when the tongue failed to contact the spout for a single cycle.

**Figure 1.**
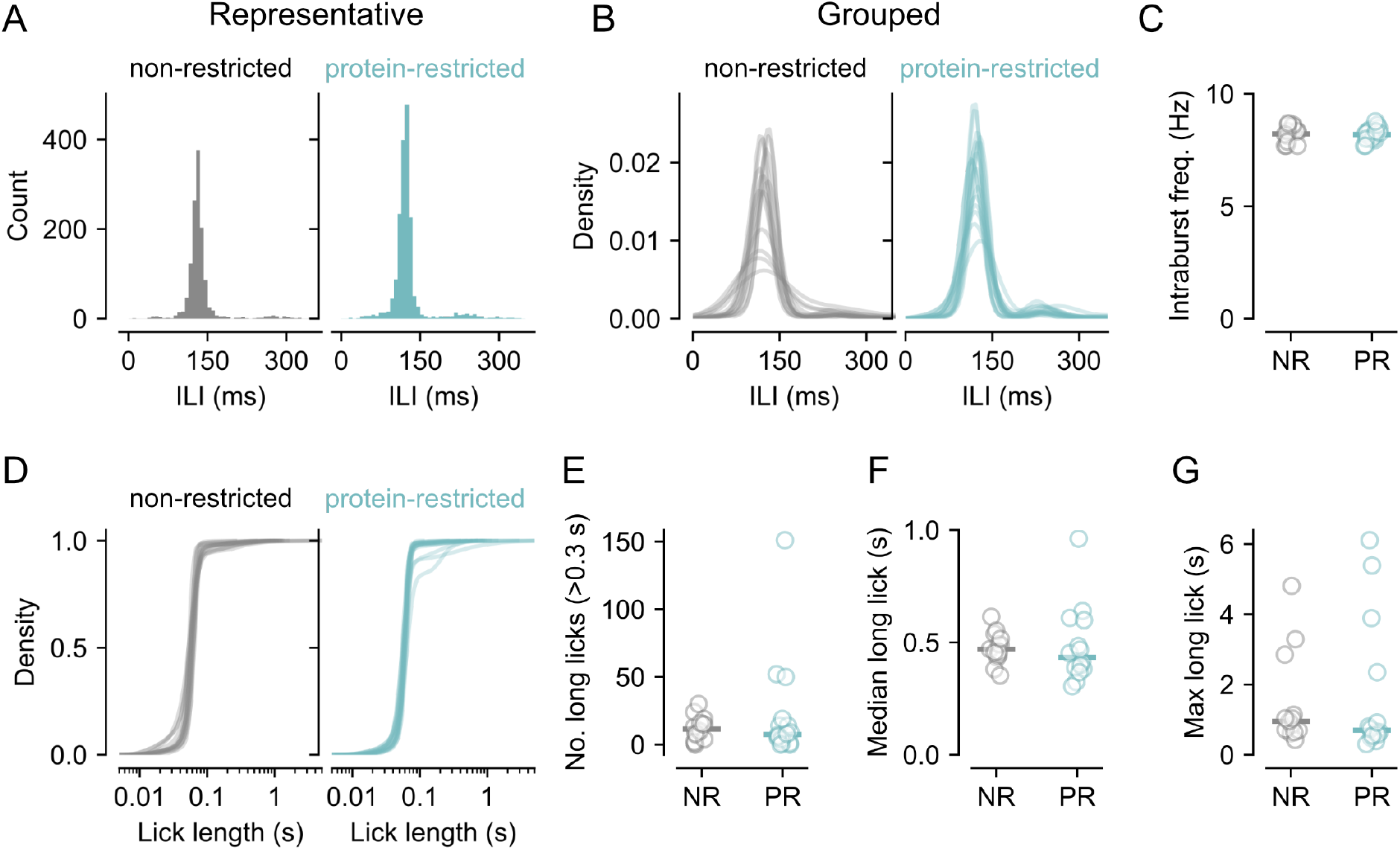
Parameters allowing quality control of lick data. **A**, Histograms showing distribution of intraburst lick intervals in representative non-restricted (NR) and protein-restricted (PR) mice. In both cases a tight distribution is seen. **B**, Kernel density estimations showing distributions in all mice overlaid. Non-restricted mice (left) and protein-restricted mice (right) show similar distributions. **C**, Mean intraburst frequency (reciprocal of mean I LI) is not different between the diet groups. Circles show individual mice and line shows mean. **D**, Cumulative density plots for lick lengths in all mice. For most mice there are very few licks that exceed 0.1 s. **E, F, G**, Number of long licks (>0.3 s), median long lick, and the maximum long lick, respectively. In no cases was there any difference between diet groups. Circles are individual mice and line shows mean.

When offsets are available, the length of each lick can be calculated and long licks can be identified (e.g., licks when the tongue remains in contact with the spout for longer than a defined threshold). Here, from the same data set, we used a threshold >300 ms to determine the number of long licks for each mouse. Inspection of cumulative lick lengths shows that almost all licks are <1 s (Fig. 1D). Most mice had very few long licks (Fig. 1E; mean: NR, 11.7 vs. PR 21.3; t = .978, p= .342) and the proportion of long licks to each mouse’s total licks was on average 0.01, or 1%, for both NR and PR mice (Fig. 1E). Moreover, the majority of long licks were <500 ms and none were longer than ∼6 s (Figs. 1F and Fig 1G). Again, there were no group differences in the average length of long licks (t = .174, p = .864) or the maximum long lick (t = .364, p = .719).

### Using microstructure to understand reasons behind a difference in intake

Following quality control of the dataset, we proceeded to analyse the classical measurements resulting from lick microstructure analysis. First, we noted that total licks for “Ensure” were higher in PR mice than NR mice (Fig. 2A; t = 3.23, p = .004) and examination of cumulative intake (Fig. 2B) suggested that the difference in licks was apparent even relatively early in the session. To examine how these licks were organised, we calculated burst number and burst size. Burst number was significantly higher in PR mice than in NR controls (Fig. 2C; t = 2.80, p = .010) but burst size did not differ by diet groups, either across the whole session (Fig. 2D; t = .739, p = .467) or when only the first 3 bursts were considered (Fig. 2E; t =1.102, p= .281).

**Figure 2.**
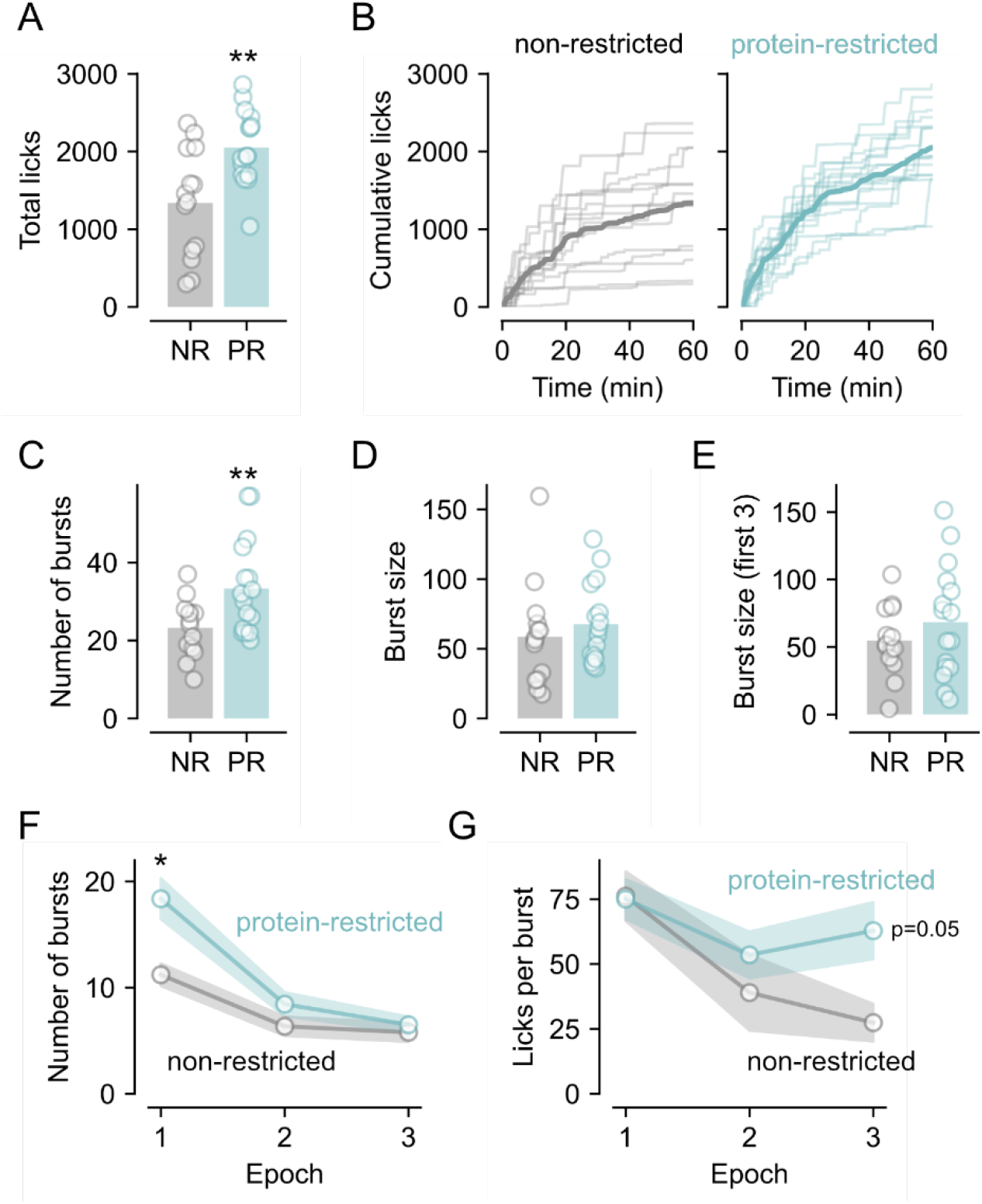
Microstructural parameters in mice consuming “Ensure”. **A**, Protein-restricted (PR) mice lick more for Ensure than non-restricted (NR) control mice. **B**, Cumulative licks for Ensure in non-restricted and protein-restricted mice. **C**, Protein-restricted mice exhibit a greater number of bursts than non-restricted mice. **D, E**, There is no difference in burst size between diet groups regardless of .whether the entire session or just the first 3 bursts are analysed. **F**, When the session is divided into thirds, number of bursts was found to be higher in protein-restricted mice in the early part of the session but this difference between groups had diminished by later in the session. **G**, With burst size, there was only a trend towards a difference between groups in the last third of the session. In A, C, D, and E circles are individual mice and bars represent mean. In F and G, circles show mean and shaded area is SEM. *, * p < 0.05, 0.01 vs non-restricted control mice.

Based on the apparent differences across time, we also used the function in *lickcalc* that allows a session to be divided into epochs. Here, we divided the session into thirds by time and analysed how burst characteristics changed across time (Figs. 2F and 2G. Repeated measures ANOVA revealed that for burst number there were main effects of epoch (F_2,56_ = 39.679, p <.001), diet group (F_1,28_ = 7.3, p = .012), and a significant epoch x diet interaction (F_2,56_ = 4.970, p = .010). Post hoc Holm tests revealed that the diet groups differed in epoch 1 (p = 0.015) but not during epoch 2 (p = .354) or epoch 3 (p = .553). For burst size, analysis revealed that there was a significant main effect of epoch (F_2,56_ = 6.853, p = .002), but no effect of diet group (F_1,28_ = 2.506, p = .125), and no interaction (F_2,56_ = 2.047, p = .139). Holm contrasts revealed that burst size did not differ in epoch 1 (p = .927) or epoch 2 (p = .810) but that there was a trend for protein-restricted mice to exhibit more licks per burst in epoch 3 (p = .051).

### Using microstructure to reveal differences other than intake

Next, we demonstrate that microstructural analysis can reveal differences between experimental groups even in the absence of an effect on total licks. Here, NR and PR mice were licking for Scandishake (a low protein version of Ensure). Analysis of total licks revealed that there was no overall difference in intake between diet groups (Figs. 3A and 3B; t = .497, p = .628). Neither was there a significant difference in number of bursts (Fig. 3C; t = 1.501, p = .161). Burst size, however, was smaller in PR mice than NR mice over the entire session (Fig. 3D; t = 2.407, p = .045) although there was no difference when only the first three bursts were considered (Fig. 3E; t = 1.227, p = .259). The interpretation of this pattern of results is that the diet groups did not differ in post-ingestive satiety and/or motivation to consume Scandishake (no difference in total licks and number of bursts) but that the PR mice did find Scandishake less palatable than NR mice (decreased burst size).

**Figure 3.**
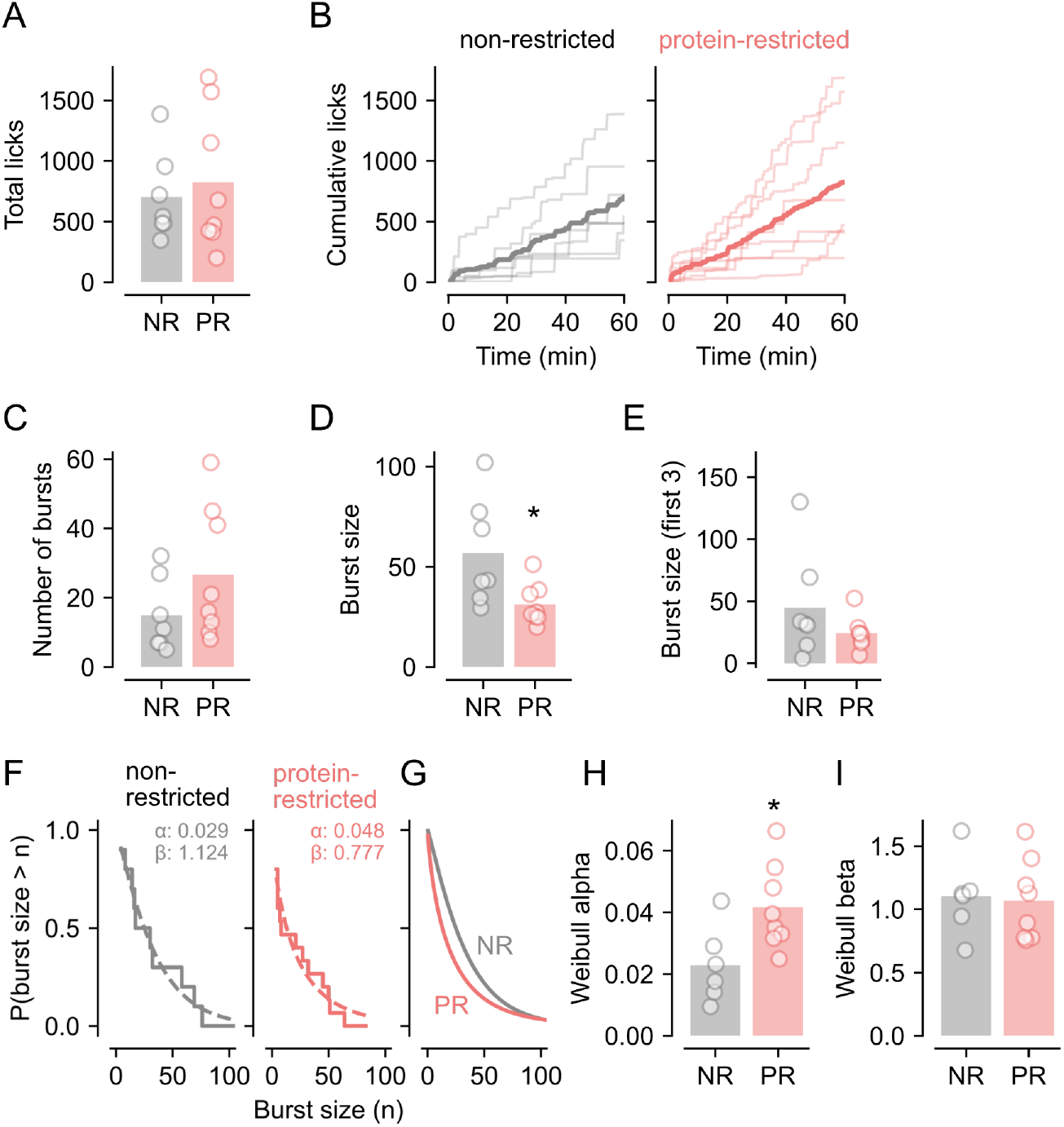
Microstructural parameters in mice licking for Scandishake (low protein Ensure). **A**, No difference in total licks between non-restricted (NR) and protein-restricted (PR) mice. **B**, Cumulative licks for Scandishake in non-restricted and protein-restricted mice. **C**, No difference in number of bursts between diet groups. **D, E**, Reduction in burst size in protein-restricted mice, relative to non-restricted mice observed across the whole session but not for the first 3 bursts. **F**, Weibull fits (dashed lines) to cumulative probability (solid lines) from representative NR and PR mice. **G**, Group averages of Weibull fits. **H**, Weibull alpha parameter is greater in PR mice than NR mice indicating more rapid decay of burst probability with increasing licks per burst. **I**, No difference in Weibull beta parameter between groups showing that an simple exponential fits group data well although there is substantial variability in the shape across individual mice. Plotting conventions as in Fig. 2. *, p < 0.05 vs. non-restricted control mice.

For analysis of these data, we also took advantage of an alternative method of quanitifying changes to the burst distribution that was proposed in Smith (1998), and that is included as an option in *lickcalc*. Here, the cumulative probability of bursts is fitted with a Weibull distribution yielding an alpha and a beta parameter, representing the decay and the shape of the distribution, respectively (Fig. 3F, G). Comparison of the derived parameters between diet groups showed that the Weibull alpha parameter was greater in PR mice than in NR mice (Fig. 3H; t = 2.69, p = .020) meaning that the cumulative probability decayed more quickly as burst size increased supporting the finding that mean burst size was lower in these mice. The Weibull beta parameter did not differ between groups (Fig. 3H; t = .21, p = .839) and on average was close to 1 meaning that average data approximated a simple exponential decay. However, the variability in this beta parameter across mice reflects the heterogeneity of licking distributions in the underlying data.

In summary, comparison if Weibull parameters offer an alternative to simply using mean burst size as this value may not be the best representation of the underlying burst distribution.

### Sensitivity to subtle changes in intraburst parameters

Most studies of lick microstructure do not consider subtle changes within bursts, both interlick intervals and lick lengths. Initial conclusions – based often on recording technology and analysis techniques with low temporal resolution – were that these parameters should be relatively stable across conditions as they are driven by a central pattern generator. Changes in lick microstructure, e.g., number and size of bursts, were determined by when this CPG was engaged. However, this view has been challenged and some have suggested that analysis of intraburst parameters might provide further insight into the control of ingestion, especially when considering how animals behave towards different solutions. Here, we compared our data with mice consuming either Ensure or Scandishake on two parameters relating to such changes, namely, the intraburst lick rate and lick lengths (contact time with the spout).

First, we compared the intraburst ILI across diet conditions and solutions (Fig. 4A). We found that that intraburst ILIs were shorter when mice were licking for Scandishake than for Ensure (main effect of Solution: F_1,41_ = 16.47, p < .001) with no effect of diet (main effect of Diet: F_1,41_ = .00, p = .952; Diet x Solution interaction: F_1,41_ = .06, p = .803).

**Figure 4.**
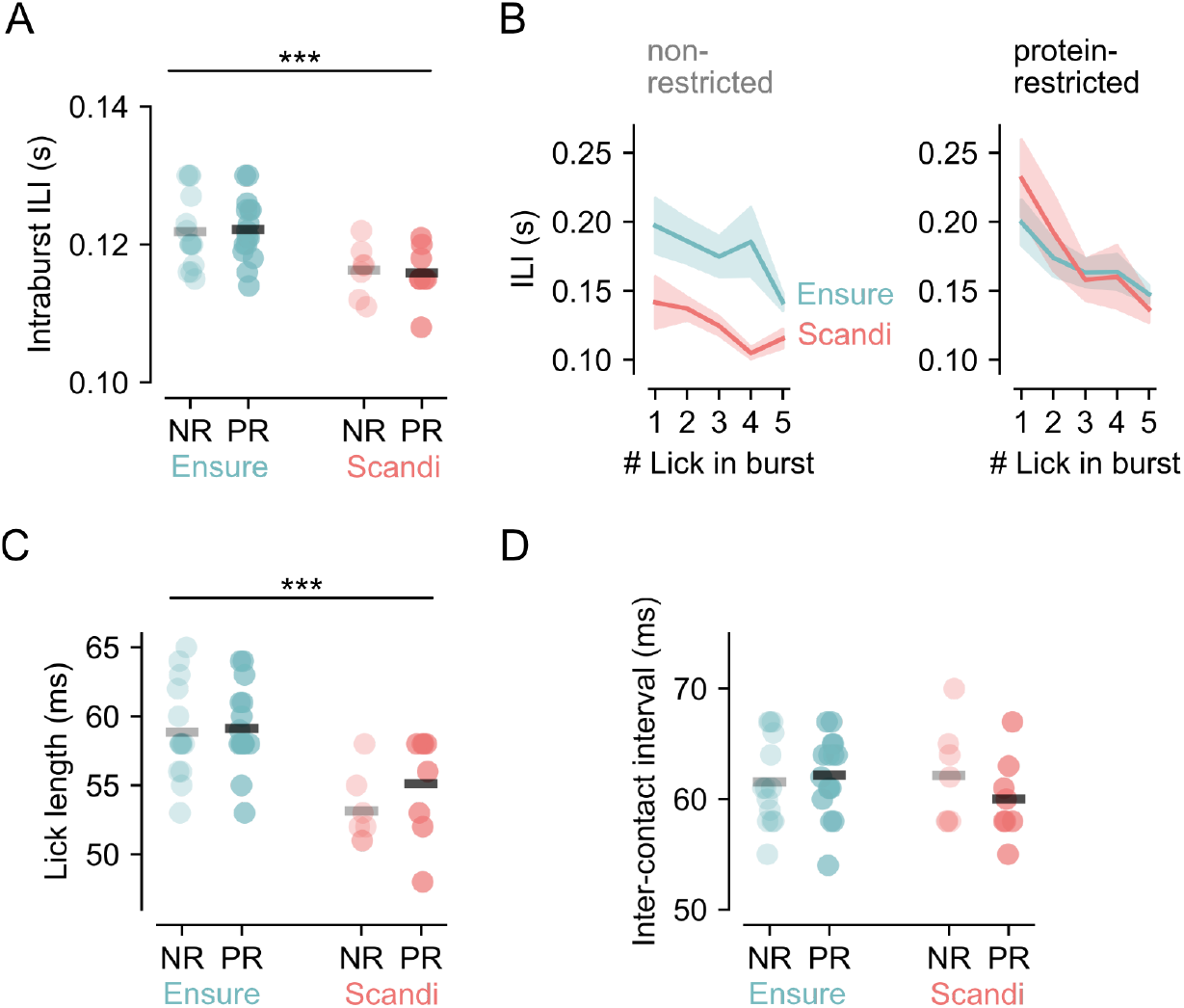
Intraburst parameters in mice licking for Ensure (high protein) and Scandishake (low protein). **A**, Mode of intraburst interlick intervals (ILIs) in non-restricted (NR) and protein-restricted (PR) mice licking for Ensure (blue) and Scandishake (Scandi; red). Intraburst ILIs are shorter when mice are licking for Scandishake than for Ensure. Circles are data from individual mice and thick lines show group mean. ***, p < 0.001 main effect of Solution. **B**, Interlick intervals (ILIs) separating first five consecutive licks per burst in NR (left) and PR (right) mice. Data are group mean (thick line) and SEM (shaded area). **C**, Mode of lick length in NR and PR mice licking for Ensure and Scandishake. Length of time tongue is in contact with the spout is shorter when mice are licking for Scandishake vs. Ensure. **D**, Mode of inter-contact intervals in NR and PR mice licking for Ensure and Scandishake. No difference in the length of time tongue is not in contact with the spout between licks. Plotting conventions for C and D as in A.

Next, we tested whether intraburst ILIs changed during a burst by examining the first five intraburst ILIs (Fig. 4B). This analysis showed that there was a highly significant effect of Lick # on ILI with ILIs decreasing throughout the burst (F_4,164_ = 12.17, p <.001). There were no interactions between Lick and Solution or Diet (all ps >.05) but there was a significant interaction between Solution and Diet (F_1,41_ = 4.91, p = .032). Post hoc Holm tests revealed that there was a trend for ILIs for Ensure and Scandishake to differ but only for NR (p = .059).

Finally, to better understand why intraburst ILIs were shorter for Scandishake vs.Ensure, we divided ILIs into the time the longue was in contact with the spout (lick length) from time between contacts (intercontact time). We found that lick length was shorter for Scandishake than for Ensure (Fig. 4C; main effect of Solution: F_1,41_ = 21.81, p < .001) with no effect of diet (main effect of Diet: F_1,41_ = .75, p = .393; Diet x Solution interaction: F_1,41_ = .69, p = .410). In contrast, the intercontact time was not affected by Solution or Diet (Fig. 4D; main effect of Solution: F_1,41_ = .54, p = .466; main effect of Diet: F_1,41_ = .07, p = .794; Diet x Solution interaction: F_1,41_ = 1.27, p = .267).

In summary, we identified subtle changes to lick microstructure within bursts that were driven by changes between solutions but relatively unaffected by dietary manipulations. Moreover, changes in ILIs resulted from differences in contact time between the tongue and the sipper, rather than in time between each lick.

## Discussion

The software presented in this manuscript, *lickcalc*, allows users to easily analyse several features of licking behaviour. This serves multiple purposes. First, data can be subjected to rigorous quality control checks to ensure hardware are set up and functioning properly and to identify common issues when using contact lickometers. Second, considering microstructure of licking behaviour goes beyond simply looking at changes in intake and provides insight into the psychological and physiological drivers of ingestive behaviour.

With respect to quality control, we show that examining the distribution of ILIs and the occurrence of long licks allows issues such as improperly positioned spouts to be identified. The ability to easily see the effect of changing parameters such as ILI threshold and minimum burst length lets researchers gain a deeper understanding of the processes involved in analysis of licking behaviour. Importantly, these parameters are noted in the outputs from *lickcalc*, which will aid in appropriate reporting of methods and reproducibility of analysis pipelines.

To highlight the utility of lick microstructure analysis, we present data describing two scenarios in which such analysis can reveal differences between experimental groups. The first addresses the reasons underlying differences in intake, while the second reveals differences between groups that are not apparent when only considering total intake. Concurrent with the first scenario, mice that are protein-restricted lick more for “Ensure” than non-restricted controls do, and they do so because they have more bursts of licking, without a change in the average number of licks in each burst. This implies that the increase in consumption when restricted is driven by a reduction in post-ingestive feedback rather than orosensory feedback. An example of the second scenario is when licking for a low-protein version of Ensure, Scandishake, there is no difference between diet groups in overall intake. Nonetheless, microstructure reveals that the pattern in which NR and PR mice is quite different, with PR mice exhibiting significantly fewer licks in each burst of licking. Our interpretation is therefore that PR mice find Scandishake less palatable than NR mice but reach equal intake despite the change in hedonic valuation of the solution. In other words, protein-restriction affects responses to nutrient solutions that are more nuanced than just changes in intake.

This software includes some analysis options that are not commonly used in lick microstructure analysis. One of these is the option to fit data with a Weibull distribution and derive alpha and beta parameters that describe the underlying distribution of burst sizes. These parameters, particularly alpha representing the decay constant, may describe the data better than the mean burst size which is commonly used and this, in turn, can provide more power and larger effect sizes. This analysis is not always appropriate, however, and is unreliable if the Weibull fit is poor (i.e., low r^2^ values). Poor fits are common when they are few bursts which can occur due to certain manipulations or when drinking unpalatable solutions. Thus, care should be taken to choose the appropriate method depending on the experimental set-up and observed data.

The software also provides detailed analysis of intraburst parameters. Most studies of lick microstructure focus on burst number and size while downplaying changes to intraburst variability. This is due to the long held belief that as high frequency licking is generated by a central pattern generator (CPG), changes to motivational states determine when the CPG is engaged without altering its output. This view has been challenged, however, and more in-depth analysis of intraburst parameters are likely warranted (X. B. Lin et al., 2013). Indeed, here we show here that intake of two different solutions results in a change to the intraburst ILI, which is explained by an increase in the time the tongue contacts the drinking spout on each lick. By providing these parameters as part of the analysis suite, we propose that more researchers should consider intraburst effects as core microstructural outputs.

Our software accepts several file formats including basic CSV/TXT files, Med Associates (columnar or array-like), Coulbourn Instruments, and OHRBETS (Gordon-Fennell et al., 2023). Adapting the software to accept a format not on this list is possible either by forking the repository and adding a new file parser or by submitting an issue on the project’s Github site. Previous custom requests for new lab-specific file parsers have been handled in 1-2 days.

Using the “long lick” functionality requires both the onset and offset of each lick to have been recorded. Unfortunately, many systems do not measure lick offsets, at least as default behaviour. For those using Med Associates who want to record offsets, a template MPC file is provided in the repository with instructions (https://github.com/uit-no/lickcalc/blob/master/assets/examples/offsets_template.MPC). Users of other systems could adapt the logic contained within this file to their own setups.

*lickcalc* allows for a high degree of customisation via loading of an optional configuration file. The config file contains settings for each of the key parameters and even allows users to override default values on plots and sliders. One user has reported using this option to perform analysis of meal pattering (e.g., meals per hour) rather than lick microstructure. Beyond this, as open source software, the codebase itself is available on Github and requests for new features are welcomed.

In conclusion, *lickcalc* makes analysis of lick microstructure straightforward for all researchers and, as browser-based software, does not require custom coding or even a local installation. We hope this will encourage more researchers to probe their datasets more thoroughly as well as adding consistency and transparency in how results from lick microstructure experiments are conducted and reported.

## Acknowledgements

We acknowledge contributions from colleagues in the field of ingestive behaviour who have thought deeply about the meaning of patterns of licking. In particular, the following have either contributed data for us to test or have advised on the design of the program and analysis: (in alphabetical order) Derek Daniels, Samantha Fortin, Adam Gordon-Fennell, Kevin Myers, Jess Santollo, Lindsey Schier, and Alan Spector. In addition, we acknowledge help and advice provided by UiT’s section for digital research resources and infrastructure, especially Rolf Andersen.

## Notes

**Conflict of Interest:** Authors report no conflict of interest.

https://github.com/mccutcheonlab/lickcalc-paper

### Competing Interest Statement

The authors have declared no competing interest.

### Summary of Updates

Updated functions in software described in manuscript and more infortmation on interpretation of microstructure properties.

